# Dorsal horn dendrite polarization during Spinal Cord Stimulation (SCS) predicted using the quasi-uniform-mirror assumption

**DOI:** 10.1101/2024.01.17.575924

**Authors:** Adantchede Louis Zannou, Mojtaba Belali Koochesfahani, Marc Russo, Marom Bikson

## Abstract

While the mechanisms of Spinal Cord Stimulation are traditionally assumed to follow pacing of dorsal columns axons, the development of sub-paresthesia SCS and new waveforms has encouraged consideration of different cellular targets. Given their relative proximity to the stimulation electrodes and role in pain processing (e.g., synaptic processing and gate control theory), spinal cord dorsal horn interneurons may be direct targets of stimulation. We developed a novel computational modeling pipeline termed ‘quasi-uniform-mirror assumption’ and apply it to predict polarization of dorsal horn inter-neuron cell types (Islet type, Central type, Stellate/Radial, Vertical-like) to SCS. The quasi-uniform-mirror assumption allows prediction, for each cell type and location in the dorsal horn, the peak and axis of dendritic polarization as well as the impact of stimulation pulse width. For long pulse (dc), the peak polarization per mA of SCS with a bipolar montage were: Islet type 3.5 mV, Central type 2 mV, Stellate/Radial 1.4 mV, Vertical-like 2.5 mV. For Stellate/Radial the peak dendritic polarization was dorsal-ventral, and for Islet type Central peak dendrite polarization was in the rostral-caudal axis. For Islet type and Central type cells peak dendrite polarization was between stimulation electrodes, while for Stellate/Radial and Vertical-like peak dendrite polarization was under stimulation electrodes. The impact of pulse width depended on cell membrane time constants, which in the quasi-uniform-mirror assumption are uniform first-order. Assuming 1 ms time constant, for a 1 ms or 100 us pulse width, the peak dendrite polarization decreases (from the dc values) by ∼33% and ∼88%. These results suggest, for the conditions modeled, maximum polarization is of islet-cells in the superficial dorsal horn at locations between electrodes; applying 2 mA 1 ms pulse SCS, the peak islet-cell dendrite polarization is ∼4.7 mV. Polarizations of a few mV have been shown to modulate synaptic processing through sub-threshold mechanisms.

## Introduction

The canonical theory of pain reduction by neuromodulation is gate control [1] involving synaptic processing by spinal cord dorsal horn interneurons. Spinal Cord Stimulation (SCS) is often explained by direct stimulation of dorsal column axons leading to indirect activation of dorsal horn interneurons [2], [3]. The traditional focus on dorsal column stimulation is for several reasons. One, other forms of pain neuromodulation, such a dorsal root ganglion or peripheral nerve stimulation, evidently effects axons. However, SCS may work differently. Two, the electric fields produced in the dorsal horn are reduced compared to in dorsal columns [4], [5], presumably below action potential activation threshold. However, sub-threshold stimulation mechanisms by low-intensity electric fields are established including through polarization of dendrites and changes in synaptic processing [5]–[7]. Thus, sub-threshold stimulation may modulate pain gate related synapse. Third, paresthesia, and therefore dorsal column stimulation, was considered require for pain relief by SCS [8]. New SCS paradigms show this is not the case [2]. Moreover, many of these novel waveforms are surprising based on dorsal column pacing hypotheses [9]–[11], and some SCS approaches are now designed for dorsal horn stimulation [12]. There is thus a re-evaluation of the role of direct dorsal horn interneuron stimulation in SCS [5], [13]–[15] including islet cells.

There is long standing research showing synaptic currents generated in the dorsal horn can be recorded epidurally [16], [17]. We showed synaptic signals from the dorsal horn can be evoked and recorded with epidural leads; these signals termed ‘eSAPS’ are distinct from ‘eCAPS’ of dorsal column axon origin [18]. Synaptic activity in the dorsal horn is of interest for its relation to gate control mechanisms. Based on the principle of reciprocity [19]–[21], the recording of signals of synaptic-origin infers the ability to stimulate the same processes (e.g., dendrites) generating those signals.

Computational models inform explanations for and the design of SCS. Models suggest what cells and what compartments of those cells are activated by SCS. Analysis focused on supra-threshold (e.g., pacing) mechanisms has traditional focused on axons [22], [23]. However, in analysis of sub-threshold modulation the axon terminal/synapse [5], [7], [24], dendrite [6], [15],, and soma [26] compartment polarization are shown to influence brain processing. Here, we are interested in the question: how much are dorsal horn interneurons polarized by SCS. Which neurons are polarized and how much determines the mechanisms of SCS and so its design. Alongside computational approaches simulating intricate neuron and network details [15], [27], there have been developments towards “semi-analytic” models with reduced complexity [28]–[30], [30], [31]. To this end, two previously developed simplifications, the *mirror estimate* [32] and the *quasi-uniform assumption* [33]–[35], are unified here for the first time. Given an SCS dose (current, electrode configuration, pulse duration) our *quasi-uniform-mirror* approach yields predictions of peak polarization, at dendrite terminals, across prototypical types of dorsal horn interneurons.

## Methods

### General Approach and Quasi-Uniform-Mirror Rationale

The novel quasi-uniform-mirror assumption models derives from the following assumptions:

1. The conventional *quasi-static assumption* in neuromodulation regarding the physics of current passage through macroscopic tissue [36]–[38], including SCS [39], leading to the governing Laplace equation. The electric field generated in the spinal cord by SCS is a function of the stimulation dose (electrode position, current) and the anatomy/macroscopic tissue properties [40].
2. The *quasi-uniform assumption* [34], [35] where the membrane polarization of each neuron is well approximated by assuming a uniform electric field across the neuron (i.e., each neuron can be modeled exposed to a single electric field vector). For our purposes, we decompose the electric field into Dorsal-Ventral, Rostral-Caudal, and Medial-Lateral electric field components. One aspect of the quasi-uniform assumption is that animal models either implicitly rely on it for translation of dose or explicitly apply uniform electric fields – this includes stimulation of spinal cord slices [16].
3. The *mirror estimate* assumption [32] where the membrane polarization of each given neuron compartment is approximated by the average voltage outside all compartment minus the outside voltage of the given compartment. The mirror estimate follows from assuming a neuron is electronically compact. The mirror estimate allows for ignoring membrane properties, as static conditions are considered, and it is more accurate with increasing pulse width. While the mirror estimate can be applied for all neuron compartments, for our purposes, for each neuron we will predict the maximum membrane polarization (that occurs at dendrite process ends).
4. We assume that the density of compartments per volume is equal across the entire extent of the neuron’s dendrite. We ignore the contribution (presence) of the axon. For the purpose of our *quasi-uniform-mirror* pipeline this allow two further approximations:
  a. To represent a neuronal morphology by its maximal dendrite distance in the Dorsal-Ventral, Rostral-Caudal, and Medial-Lateral directions. Based on aggerate literature, each cell (e.g., “islet-type” cells) is assigned three dimensional values.
  b. That the peak polarization profile along any axis in cell morphology will be a linear transition from peak hyperpolarization at one to peak depolarization at the other end. The peak polarization along each axis (which will be at the associated dendrite ends) is equal to half of the magnitude the electric field in the axis times the distance from end to end for that axis. Since this combines 2 and 3, we call this the *quasi-uniform mirror assumption*.
5. In addition to assigning each cell type three morphological dimension values we further assume each neuron type fills the entire region of interest: the spinal cord grey matter dorsal horn. Specifically in each point in space of the dorsal horn grey matter a neuron is represented. Combining 4 and 5, we predict the maximum polarization for a cell type centered at each point in space in the grey matter. This generates a continuous map across the dorsal horn of predicted maximum membrane polarization for a given cell type.
6. While the mirror estimate assumes a sufficient long stimulation pulse width (relative to membrane time constant) for membrane compactness, we assume membrane polarization follows first order kinetics based on given time constant. For sufficiently long pulses (e.g., approaching DC) the kinetics term drops.

A result of the combination of these assumptions is that for each point in a region of interest (e.g., the grey matter of the spinal cord) for any given stimulation dose (specified by stimulation intensity, electrode position/size, and waveform) one can predict the maximum dendrite polarization of given neuron types (specified by their dendrite morphology vector and time constant) (*Equation 1*).

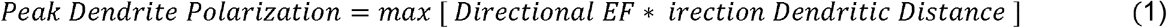

### Spinal Cord Stimulation Dose and Current Flow Model

For spinal column anatomy we adapted a previously developed models [40], [41]. The geometry was computer-aided design (CAD) of the of lower thoracic (T8–T12) spinal cord with eight tissue compartments, including the vertebrae, intervertebral disc, soft tissues, epidural fat, dura, cerebrospinal fluid (CSF), white matter and grey matter in SolidWorks 2016 (Dassault Systemes Americas Corp, Waltham, MA). A simulated SCS lead (electrode diameter: 1.3 mm; electrode length: 3.0 mm; edge-to-edge interelectrode spacing: 4 mm) was positioned in the epidural fat along the mediolateral midline of the spine. The entire CAD model assembly was then segmented and meshed into a finer mesh using a built-in voxel-based adaptive meshing algorithm of Simpleware ScanIP (Synopsys Inc, Mountain View, CA). Mesh density was refined until additional model refinement produced a <1% difference in peak temperature change and peak electric field. The resulting mesh consisted of >5.6 million tetrahedral elements. The final FEM model was later imported and computationally solved in COMSOL (COMSOL Multiphysics, Boston, MA). We assumed a quasi-static electrical conduction model, which allowed us to apply Laplace’s equation to compute the electric potential V (Volts). In most analysis we normalize results to ‘per mA’ as the assumptions on current flow and polarization developed here are linear.

### Interneuron Classification, Morphological Parameters, and Time Constants

There is ongoing discussion about how to classify distinct interneuron types in the spinal cord dorsal horn based on morphological, biophysical, generic, or functional methods. There have been different names proposed as well as the suggestion of a spectrum, rather than distinct categories, or cells. For our purposes, it was sufficient to identify four prototypical dorsal horn interneuron types, that are commonly recognized (across species, even if with different names) of distinct dendritic morphology and functions. For each type, we select a maximal dendrite distance in the Dorsal-Ventral, Rostral-Caudal, and Medial-Lateral directions (Table 1). We do not consider axons (when identified) in these dimensions, and generally aim for a median value across studies. A virtue of our pipeline is it can be easily adapted to different cell types or morphologies. We emphasize features within individual cell types vary, and even the nature of classification is debated, but for our purposes the goal is to identify canonical and distinct cell archetypes – and for this reason we refer to each class as “type”.

**Table 1:** Cell types and their dendrite distances. Values are representative dendrite distances across indicated references.

| Cell Type | Dorsal-Ventral | Rostral-Caudal | Medial-Lateral | References |
| --- | --- | --- | --- | --- |
| Islet Cell Types | 117 $\mu\text{m}$ | 642 $\mu\text{m}$ | 99 $\mu\text{m}$ | [42]–[48] |
| Central Cell Types | 138 $\mu\text{m}$ | 327 $\mu\text{m}$ | 210 $\mu\text{m}$ | [43], [44], [46], [47] |
| Stellate/Radial | 189 $\mu\text{m}$ | 244 $\mu\text{m}$ | 206 $\mu\text{m}$ | [42]–[44], [46], [47] |
| Vertical-Like | 247 $\mu\text{m}$ | 293 $\mu\text{m}$ | 259 $\mu\text{m}$ | [42], [44], [46], [47] |

Time constants for polarization are challenging to infer. Moreover, our pipeline assumes a dendrite axis-wide first-order time constant. Distinct time constants are expected between cell types and across compartments (soma vs axon vs dendrite), and will be a function of dynamic membrane conditions (e.g., synaptic input). Experiential data on dendritic terminal time constants for extracellular stimulation are limited. For example, in quiescent hippocampal neurons, 8 ms dendritic time constants have been shown [6] or chronaxies of 1 ms [49]. Time constants for electrical stimulation may be inferred from strength duration curves (e.g., assuming a chronaxie = 0.7 x time constant) but depend on the stimulation method and reflect supra-threshold outcomes. Inferring time constants from simulations of complex neuronal models, it itself highly sensitivity to membrane/ion/activity parameters selected. A wide range of time constants have thus been suggested under varied assumptions and conditions [50], [51]. For our analysis we consider a range of time constants (1-16 ms).

## Results

Our modeling pipeline (Figure 1) is implemented in three steps: 1) simulation of electric fields in the tissue, specifically directional electric fields in the dorsal horn; 2) for a long (dc) pulse, prediction of maximum dorsal horn interneuron dendrite polarization by cell type; 3) further modeling of dendrite polarization considering varied pulse width.

**Figure 1.**
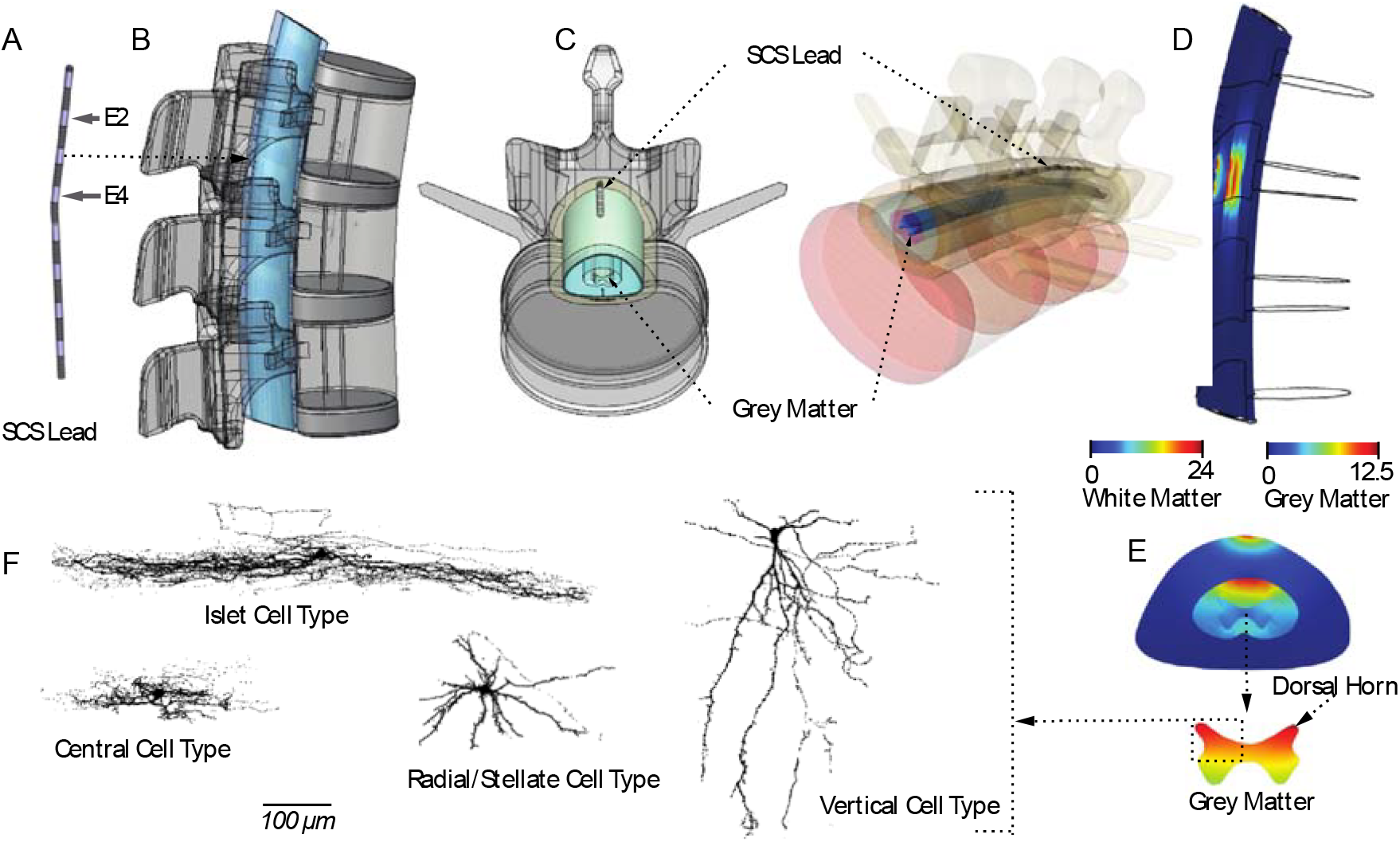
Novel quasi-uniform-mirror modeling pipelines applied to SCS of dorsal horn interneurons, A. Commercial eight-contact SCS lead is simulated with a bipolar (E2, E4) electrode montage. B, C and D. CAD–derived human spinal cord anatomy of the lower thoracic spine (T8– T10) with the eight-contact SCS lead implanted within the epidural fat. E and F. FEM simulation results predicting electric field across the spinal tissues shown transversally (E) and cross-sectionally (F). Note the electric field is reduced in the grey matter compared to white matter so we represent the electric field in each region on its own scale. G. Four different cell types of the dorsal horn are identified: Islet cell type, Central cell type, Radial/Stellate cell type, Vertical cell types. Cell types are based on a literature review. with the specific cell morphologies shown adapted from Yasaka *et al* 2010. Only the dorsal-ventral and medio-lateral dendritic structure is considered relevant for our results.

SCS produces electric fields in the spinal cord dorsal columns and grey matter with magnitude and spatial distribution consistent with prior studies (Figure 2, row 1). For the bipolar montage considered, peak electric field in the white matter are ∼24 V/m per mA, and in the grey matter ∼12.5 V/m per mA. We note while electric fields are reduced in the grey vs white matter (Figure 2, row 2), this is only by a factor of ∼2, and may even be more comparable depending in the electrode montage (e.g., inter-electrode distance) or based on tissue assumptions (e.g., grey matter conductivity. We decompose the electric fields in the grey matter into its medio-lateral, dorsal-ventral, and rostral-caudal components (Figure 2, row 3). Note between electrodes the rostral-caudal component is maximized while under the electrodes the dorsal-ventral component is maximized. The inset identifies the dorsal horn considered in the remaining analysis, only one side is considered because of lateral symmetry.

**Figure 2.**
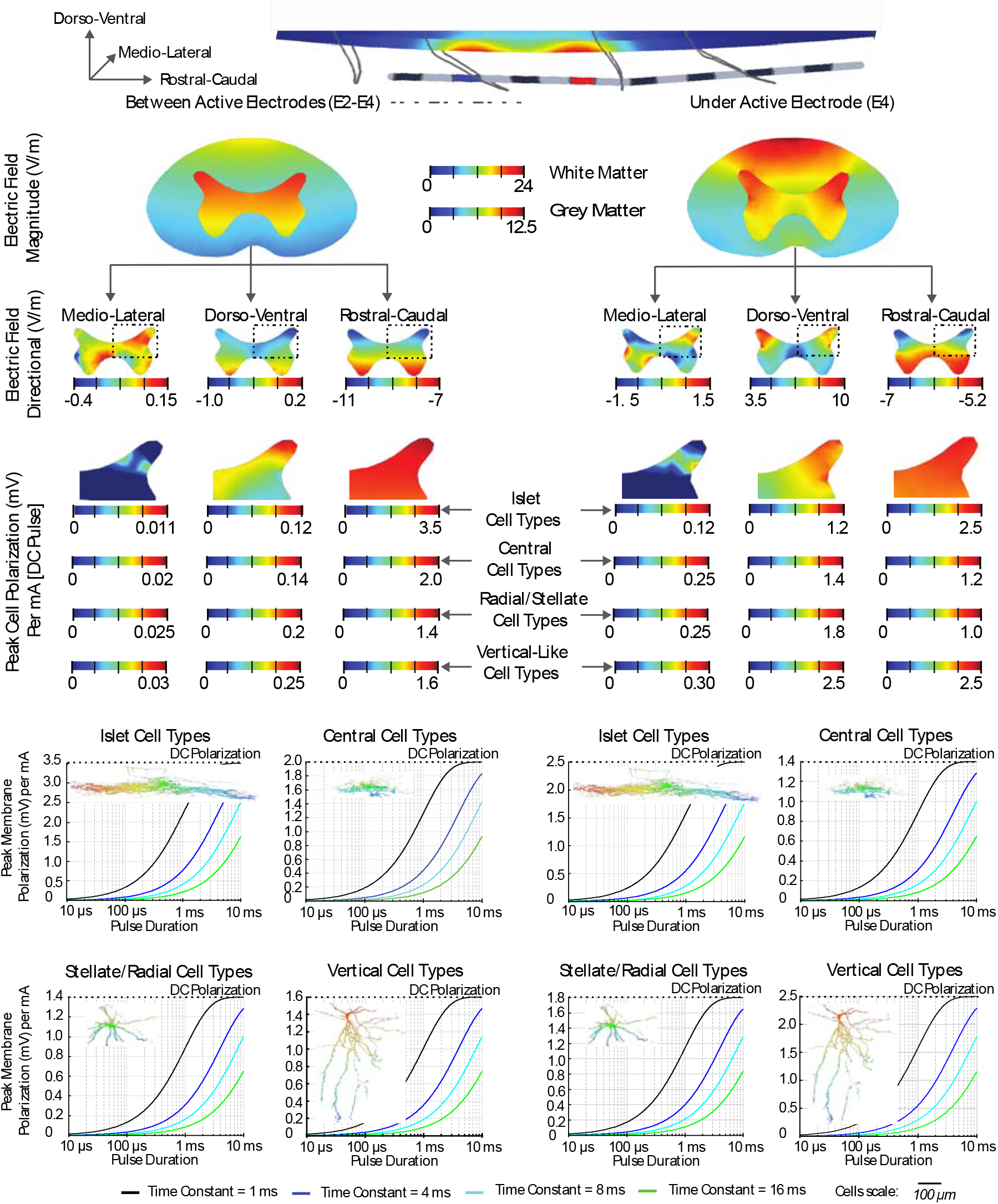
Prediction of dorsal horn interneuron polarization during SCS using a novel quasi-uniform-mirror modeling pipelines. We consider spinal cord electric field and cell polarization analysis under active electrode, E4 (right columns) and between active electrodes, E2-E4 (Left columns). Rows 1 and 2. Magnitude of electric fields in the grey matter and white matter of the spinal cord. Row 3. Directional analysis (medio-lateral, dorsal-ventral, and rostral-caudal components) of electric fields in the grey matter. Row 4. Directional peak cell polarization analysis across Islet, Central, Radial/Stellate, and Vertical neural cell types. Rows 5 and 6. Peak cell membrane polarization evaluation at various stimulation pulse duration assuming membrane polarization time constants of 1 ms, 4 ms, 8 ms, and 16 ms across Islet, Central, Radial/Stellate, and Vertical neural cell types. Insets indicate simulated Islet, Central, Radial/Stellate, and Vertical neural cell polarizations. The false color map is a relative scale with peak dendrite depolarization and hyper-polarization opposite ends (red and blue) with a peak amplitude corresponding to any associated stimulation current, pulse width, and membrane time constant.

The dendrite polarization, but cell type, is predicted, first considering a long (dc) pulse (Figure 2, middle). The peak polarization will vary across the dorsal horn (Figure 2, row 4); this *relative* polarization does not vary by cell type so is indicated once. The absolute magnitude of the polarization will vary by cell type as indicated. The maximum polarization of 3.5 mV per mA is shown for Islet-type cells, located between the electrodes, as influenced by electric fields in the rostra-caudal direction.

The polarization given a limited duration pulse width (10 us to 10 ms), given a cell time constant (1 ms, 4 ms, 8 ms,16 ms) is predicted for each cell type (Figure 2 bottom). The maximum polarization for a dc pulse is shown (to dashed line) for each condition per mA applied, with this pulse duration-limited polarization decreasing with decreased pulse and cell time constant. As expected for a first order time constant, when the pulse width equaled the time constant, the polarization decreased by 33% from the dc value. For example, for Islet type cells located between electrodes, given a 1 ms time constant and a 1 ms stimulation pulse, peak dendrite polarization was 2.3 mV per mA applied.

Also plotted (Figure 2 bottom) is the relative polarization profile across the cell dendrite (false color). According to the quasi-uniform-mirror assumptions (see Methods) the polarization is a linear gradient from a most depolarized end to a most hyper-polarized end, with zero polarizing in the middle. The direction of the gradient is based on the electric field direction and dendrite length producing the most polarization; and so these plots ignore polarization in any other axis or from any other mechanisms. The scale of false color map of polarization is relative to the peak polarization at the distal dendrites, as given by each cells overlaid chart. For example, for an Islet-type cell located between electrodes, given a 1 ms time constant and a 1 ms stimulation pulse at 2 mA, the peak dendrite polarization is 4.6 mV (corresponding to blue and red). Note our analysis of polarization (Figure 2 middle, bottom) is agnostic to pulse polarity.

## Discussion

Notwithstanding decades of technical developments including new waveforms, positive clinical trials and expanding indications, and expanding basic science, questions remain about the mechanisms of SCS. Across approaches and theories, they remains a convergence around modulation of pain processing through the spinal cord: specifical synaptic processing by interneurons in the superficial dorsal horn, such as gate control theory [52]. With this ultimate goal of changing synaptic processing by dorsal horn interneurons, there are different indirect and direct targets for SCS. Indirect targets include stimulation of dorsal column axons whose collateral than synapse onto dorsal horn interneurons (classical SCS theory; [53]), activation of glia, or action of blood vessels (endothelial cells) inside or leading to the dorsal horn leading to secondary change in dorsal horn interneuron metabolism. The direct target would be stimulation dorsal horn interneurons directly [5], [13]–[15].

An essential consideration on the mechanisms of neuromodulation is “which neuronal elements” are excited [50], [51]); by which is meant not only what cell (types) are activated but also which structure of the cells. The considered neuronal strictures are the soma [26], dendrites [6], [54], and axon – hillock [55], branches [56], or terminal [24]. For SCS when *a priori* assuming dorsal column stimulation, axons are the neuronal element of interest. For SCS based on direct dorsal horn stimulation, all neuronal elements are present. Here we consider SCS polarization of dendrites in the spinal cord grey matter.

The attention to axons as stimulation targets derives from their low threshold to action potential generation, when the outcomes of stimulation are assumed to derive from pacing neurons (at rest). There are many applications where such analysis seems comprehensive, e.g., where the stimulation electrode is adjacent to an axon bundle (nerve). The relevance of dendrite stimulation [57]–[59] derives from several factors including their 1) essential role in all brain processing; 2) their comparable sensitivity to polarization as axons; 3) recognition that polarization below action potential threshold (sub-threshold stimulation) effects brain function.

How SCS might modulate pain through polarization of dorsal horn neurons dendrites is multifaceted, but a first step is to consider how much dendritic polarization is produced across distinct spinal interneurons. The location relative to SCS electrodes and the role of pulse duration provides further insight and links to SCS dose (waveform) optimization. These questions are addressed here though the development of a novel quasi-uniform-mirror modeling process (see Methods).

The quasi-uniform-mirror modeling depends on several explicit assumptions. These assumptions are supported by biophysical considerations developed separately for the mirror estimate (including neuronal compactness) and the quasi-uniform assumption (including limited changes in electric fields over the length of neurons). The quasi-uniform-mirror combines these thorough additional simplifying assumptions. Evidently, the accuracy of the quasi-uniform-mirror models will depend on the neuromodulation dose and targeted neurons, but it is important to recognize: 1) the types of polarization pattern predicted by the quasi-uniform-mirror matches experiments [26], [54]; and 2) models that explicitly represent hundreds of neuronal compartments each with hundreds of membrane parameters are also sensitive to assumption, but in a more complex way. While the quasi-uniform-mirror is simple and transparent how to model parameters (e.g., length, time constant) impacts results (e.g., Equation 1).

In addition to supporting heuristic considerations of SCS mechanisms, the quasi-uniform-mirror modeling has translational implications. As the entire modeling pipeline is linear, it lends itself to closed-form dose optimization (i.e., where a target is specified and an optimal SCS dose predicted; [60] and/or rapid calculations for device programmers or closed-loop applications [61]. These can include targeting specific cell type in the dorsal horn such as islet cells. The linearity also reflects the fundamental limitations of the quasi-uniform-mirror modeling.

The main result of this study is that SCS polarizes the dendrites of dorsal horn interneurons by a few mV per mA of stimulation, with the degree of polarization dependent on the neuron location relative to the stimulation electrode, neuron type, and neuron membrane time constant compared to the stimulation pulse duration. For Islet-type interneurons located between stimulation electrodes, with pulse widths as long as the membrane time constant (e.g., 1 ms) the polarization is a high as 2.3 mV per mA (e.g., ∼7 mV for 3 mA intensity). The duty cycle of stimulation will determine the relative time in this polarized state. Previous studies established that dendrite polarization of a few mV is established to modulate synaptic processing [6], [62], synaptic plasticity [63]–[65], and oscillations [66], [67]. Our results show that the direct polarization of interneuron dendrites during SCS is of sufficient magnitude to modulate processing.

## Notes

### Competing Interest Statement

MR is the President of the International Neuromodulation Society and Director at Large of the Neuromodulation Society of Australia and New Zealand. MR discloses research activities (paid to research institution) for Boston Scientific, Mainstay Medical, Medtronic, Nevro, Presidio Medical and Saluda Medical, historical equity interest in Stimwave, Lungpacer and SPR Therapeutics, and stock options in Saluda Medical and Presidio Medical. The City University of New York holds patents on brain stimulation with MB as inventor. MB has equity in Soterix Medical Inc. MB consults, received grants, assigned inventions, and/or served on the SAB of SafeToddles, Boston Scientific, GlaxoSmithKline, Biovisics, Mecta, Lumenis, Halo Neuroscience, Google-X, i-Lumen, Humm, Allergan (Abbvie), Apple, Ybrain, Ceragem, Remz. MB is supported by grants from Harold Shames and the National Institutes of Health: NIH-NIDA UG3DA048502, NIH-NIGMS T34 GM137858, NIH-NINDS R01 NS112996, NIH-NINDS R01 NS101362, and NIH-G-RISE T32GM136499.

